# GERMIN3 plays a role in plasmodesmatal gating to regulate meristem activation related to tuberisation, tuber dormancy release and stem branching in potato

**DOI:** 10.1101/2024.09.28.615581

**Authors:** Raymond Campbell, Graham Cowan, Bernhard Wurzinger, Laurence J.M. Ducreux, Jimmy Dessoly, Wenbin Guo, Runxuan Zhang, Jenny A. Morris, Pete Hedley, Vanessa Wahl, Mark A. Taylor, Robert D. Hancock

**Affiliations:** Cellular and Molecular Sciences, The James Hutton Institute, Invergowrie, Dundee, DD2 5DA, UK; Department of Functional and Evolutionary Ecology, University of Vienna, Austria; Department of Applied Genetics and Cell Biology, BOKU University Vienna; Information and Computational Sciences, The James Hutton Institute, Invergowrie, Dundee, DD2 5DA, UK

**Keywords:** GERMIN-like proteins, meristems, plant development, plasmodesmata, Solanum tuberosum (potato), tuberisation

## Abstract

- *GERMIN3* has previously been identified as a target of the tuberigen activation complex suggesting a function in potato tuberisation but its role is presently unknown.
- To understand the role of GERMIN3 we analysed morphological, agronomic and molecular phenotypes in transgenic lines.
- *GERMIN3* over-expressing lines of *Solanum tuberosum* ssp. *andigena* exhibited increased tuber yields under permissive conditions and enhanced tuber numbers. Post-harvest tuber sprouting exhibited reduced apical dominance with increased numbers of sprouts. Apical dominance was reduced in aerial tissues of mature plants where stem growth from axillary buds was activated. Similar results were observed in the commercial cultivar Desiree. Over-expression of *GERMIN3* had no impact on the expression of *SP6A*, a positive regulator of tuberisation or *TFL1B*, a negative regulator. The GERMIN3 protein localised to the endoplasmic reticulum and transient expression in *N. benthamiana* leaves resulted in plasmodesmatal gating allowing intercellular transport of GFP-tagged sporamin independent of GERMIN3 oxalate oxidase activity.
- GERMIN3 affects tuberisation and other developmental processes by facilitating meristem activation. This identifies GERMIN3 as a novel protein associated with control of plasmodesmatal transport and supports the importance of plasmodesmatal gating in the regulation of key potato developmental processes.

## Introduction

GERMINs and GERMIN-like proteins (GLPs) are water-soluble glycoproteins widely found in plants (Dunwell *et al*., 2008). GLPs belong to large gene families, often found as multiple copies at specific chromosomal loci (Ilyas *et al*., 2020; Zaynab *et al*., 2022). GERMINs and GLPs have been implicated in many biological processes including roles in signalling cascades associated with responses to both biotic and abiotic stresses (Berna and Bernier, 1999; Davidson *et al*., 2009; Gangadhar *et al*., 2021). This is in part mediated via the fact many of them have oxalate oxidase or superoxide dismutase activity (Lane *et al*., 1993; Woo *et al*., 2000) and are reported to participate in plant responses involving hydrogen peroxide (H_2_O_2_) production. In potato, a GLP is associated with heat tolerance based on the enhanced *in vitro* growth performance of over-expressing transgenic lines subjected to heat stress scenarios (Gangadhar *et al*., 2021). This tolerance was attributed to activation of transcripts encoding antioxidant enzymes and stress responsive heat shock proteins caused by enhanced levels of H_2_O_2_ in the over-expressing lines. In Arabidopsis, other GLP proteins are located in the plasmodesmata where they appear to have a role in the control of cell-to-cell trafficking (Ham *et al*., 2012).

During potato tuber development, prior to tuber induction, phloem unloading in elongating stolons is apoplastic both in the region of the apical meristem and in the subapical region (Viola *et al*., 2001). As the sub-apex swells during early tuberisation, induction of symplastic unloading (via plasmodesmata (PD)) allows increased photoassimilate delivery, and division and expansion of parenchyma cells to create the tuber. Initiation of tuber swelling involves localised changes in sugar-responsive gene expression as well as PD and phloem function (Viola *et al*. 2001), potentially allowing the unloading of phloem-mobile signalling molecules, but the triggers for, and orchestration of, these events is not currently understood.

Throughout tuber development, axillary meristems remain symplastically isolated, maintaining a tightly controlled cellular domain around the bud as dormancy is established, limiting substrate availability and necessitating the active transport of signalling compounds into the meristematic tissue. In sprouting tubers, dormancy break is associated with symplastic re-connection of the axillary meristem to the tuber phloem network allowing the free diffusion of phloem-mobile signalling molecules and stored photoassimilates into the meristematic region (Viola *et al*., 2007). These observations strongly suggest that symplastic connectivity or isolation is a key mechanism controlling stolon/tuber bud meristem activity by regulating the supply of substrates and signalling compounds. Control of symplastic continuity is regulated by PD which connect cells and tissues into symplastic domains. Despite their importance, the molecular properties of PD are largely elusive or speculative (Han *et al*., 2019), but these structures are a likely control point for tissue function and tuber life-cycle regulation.

A complex termed the tuberigen activation complex (TAC) comprising SP6A, 14-3-3s and FLOWERING LOCUS D–LIKE 1 has recently been identified (Teo *et al*., 2017). The TAC has a regulatory role in tuber initiation acting in an analogous manner to the florigen activation complex for flowering. We have recently identified a *TFL1* gene (designated *CEN/TFL1B*) that encodes a protein thought to compete with SP6A in the TAC. Binding of TFL1B in this complex results in an inactive TAC complex (Zhang *et al*., 2020), thus RNAi-*TFL1B* lines tuberise earlier than wild-type (WT) and *35S::TFL1B* lines have delayed tuberisation.

In order to identify potential TAC targets, we carried out transcriptomic analysis of non-swelling stolons from WT compared with RNAi lines silenced for *TFL1B*. As the RNAi-*TFL1B* lines tuberise earlier, the non-swelling stolons from this genotype are primed for tuberisation and so a comparison with WT should reveal genes expressed early in tuber initiation. As well as many of the known genes which showed up-regulation at the onset of tuberisation (Hancock *et al*., 2014; Park *et al*., 2022), several transcripts encoding GERMIN proteins were identified. Transcripts annotated as *GERMIN3*, *4* and *12* are expressed at levels between 200 and 700-fold higher in the stolons from RNAi-*TFL1B* lines than in those from controls. The most strongly up-regulated *GERMIN* (*GERMIN3*) was directly transcriptionally regulated by the TAC complex (Zhang *et al*., 2020).

In view of the identification of *GERMIN3* as a direct target of the TAC, we wished to investigate the role of the protein in potato. We demonstrate that over-expression of *GERMIN3* is associated with accelerated tuberisation and higher tuber yield. Moreover, over-expressing lines exhibit a meristem activation phenotype with increases in post-harvest tuber sprouting and the development of more highly branched stems as axillary buds are activated. We show that the GFP-tagged GERMIN3 protein localises to the ER and regulates cell-to-cell movement of protein via the plasmodesmata.

## Materials and Methods

### Plant material and growth conditions

*Solanum tuberosum* ssp. *andigena* accession 7540 (ADG) WT plants used for transformation were propagated in 90 mm Petri dishes containing MS medium (Murashige and Skoog, 1962) supplemented with 20 g L^−1^ sucrose and 8 g L^−1^ agar at 18 ± 4°C, 16 h light, light intensity 100 μmol m^−2^ sec^−1^. Four-week-old *in vitro* sub-cultured plantlets were transferred to 12 cm pots containing compost and grown in a glasshouse under conditions of 16 h light (18°C) and 8 h dark (15°C). Light intensity ranged from 400 to 1,000 μmol m^−2^ sec^−1^. After 6 weeks plants were moved to a controlled environment growth room under conditions of 10 h light (18°C, 80% humidity) and 14 h dark (15°C, 70% humidity), light intensity 300 μmol m^−2^ sec^−1^ and watered daily.

### Generation of GERMIN3 transgenic potato lines

The binary construct used in this study was built based on the GoldenBraid system (gb.cloning.org) (Sarrion-Perdigones *et al*., 2011). Gene specific primers containing flanking *Bsm*B1 sites were used to amplify the full length *GERMIN3* coding sequence (PGSC0003DMT400046995). The PCR-purified fragments were ligated into a pUPD entry vector using the *Bsm*BI digestion–ligation reaction protocol and confirmed by Sanger sequencing. Assembly reactions with the p2x35S (GB0222) promoter and pTnos terminator (GB0037) using *Bsa*I and *Bsm*BI (New England Biolabs, Ipswich, MA, USA) as restriction enzymes in 25-cycle digestion/ligation reactions were performed essentially as described by Sarrion-Perdigones *et al*. (2011). The resulting pDGB1_alpha1:p2x35S:StGERMIN:pTnos transcriptional unit was further combined with the pDGB2:pPnos:NptII:Tnos transcriptional unit in pDGB1_omega2 to form the final construct. Final binary vectors were transformed into *Agrobacterium tumefaciens* strain LBA4404. Agrobacterium-mediated potato transformation was performed as described previously (Ducreux *et al*., 2005).

### Promoter transactivation assays

A 3 kb DNA fragment immediately upstream of the start ATG codon of *GERMIN3* was fused to the firefly luciferase (*LUC*) reporter gene and cloned into a pGWB vector backbone. The resulting vector was transformed into *Agrobacterium* strain GV3101. The CDS of group A bZIPs 2, 27 (FDL1a), 36, 35, 66, were PCR amplified from cDNA obtained from *S. tuberosum* ssp. *andigena* and cloned after the *UBI10* promoter to provide constitutive expression. The respective vectors were transformed into *Agrobacterium* strain GV3101.

Agrobacteria carrying the respective plasmids were grown overnight and diluted to achieve an OD600 nm of 0.1. The cultures were incubated until they reached an OD600 nm of 1.0. Cells were pelleted by centrifugation (3000 g, 22 °C, 17 min) and resuspended in 5 % (w/v) sucrose solution to achieve a final OD600 nm of 0.2. For the transactivation assays cultures of the *GERMIN3::LUC* containing agrobacteria and the respective bZIPs were mixed in a ratio of 1:1 and brought to a final OD600 nm of 0.2. For the promoter only control an *Agrobacterium* strain without transferable plasmid was used for mixing with the reporter construct carrying strain. The resulting agrobacteria suspensions were infiltrated into leaves of 3 different plants from 5 week old *N. benthamiana* plants. Each construct or combinations of constructs were infiltrated into individual leaves to avoid cross contamination.

Two days after infiltration a total of 12 leaf discs of 3 different plants from 3 different leaves were prepared using a 5 mm cork borer and floated on top of 100 µl of incubation solution (1/2 MS salts; 5 mM MES pH 6.0) in wells of a white, flat bottom, chimney, 96 well plate. Samples were incubated for 30 min at 22 °C in the light. Then 100 µl of assay solution (1/2 MS salts; 5 mM MES pH 6.0; 40 µM D-luciferin) were added to each sample. The plate was immediately subjected to luminescence measurement in a TECAN Spark plate reader. Light was detected between 500 and 750 nm, signal integration time was set to 2 sec per sample per measurement, measurement interval was 5 minutes, samples were kept in complete darkness at 25°C during the whole measurement period of 16 hours.

In order to perform the extended night treatment leaf discs were harvested right before the start of the regular night in the green house where the *N. benthamiana* plants were grown in a 16 hours light / 8 hours dark cycle at 25 °C. The 480 minutes measuring time point marks the onset of the extended night treatment. For the samples containing 2 % (w/v) sucrose the assay solution was supplemented with 2 % (w/v) sucrose.

### RNA extraction and qRT-PCR

Total RNA was extracted from various potato plant tissues using a RNeasy® Plant Mini Kit (Qiagen, https://www.qiagen.com/), following the manufacturer’s instructions. The first-strand cDNA templates were generated by reverse transcription using a double-primed RNA to cDNA EcoDry™ Premix kit (TaKaRa, Clontech, https://www.takarabio.com/about/our-brands/clontech). Potato elongation factor 1-alpha (EF1-α) primers were used as a normalisation control (Nicot *et al*., 2005). The expression level of StGERMIN, StCEN/TFL1B and StSP6A were determined using the StepOnePlus Real-Time PCR system (Applied Biosystems, https://www.thermofisher.com/uk/en/home/brands/applied-biosystems.html) and StepOne Software version 2.3 (Applied Biosystems). Gene-specific primers and Universal Probe Library (UPL, Roche Life Science, https://lifescience.roche.com/) probes (Supplemental table **S1**) were used at a concentration of 0.2 µM and 0.1 µM, respectively. Thermal cycling conditions were: 10 min denaturation at 95°C followed by 40 cycles of 15 sec at 94°C and 60 sec at 60°C. Relative expression levels were calculated and the primers validated using the Delta–Delta Ct method (Livak and Schmittgen, 2001).

### RNA in situ hybridisation

RNA in situ hybridization was performed as described by Gramma and Wahl (2023). Briefly, tissue samples were taken at end of the day (ED) and were immediately transferred into freshly prepared FAA fixative (formaldehyde–acetic acid–ethanol). Samples were subsequently dehydrated with ethanol and infiltrated with paraffin wax (Paraplast, Leica) using an automated vacuum-embedding system (ASP300S, Leica). After infiltration, samples were processed in an embedding centre (EG1160, Leica). Using a rotary microtome (RM2265, Leica), 8-µm sections were made and transferred to polysine-coated slides (VWR, USA). A full length *GERMIN3* probe (PGSC0003DMT400046995) was amplified from cDNA, cloned into the pGEM®-TEasy Vector (Promega), and synthesised with a DIG RNA labelling kit (Roche). Sections were scanned by the Axio Scan.Z1 slide scanner (Zeiss, Germany).

### RNA-seq and transcriptomics analysis

Total RNA was quality checked using a Bioanalyzer 2100 (Agilent) and sequencing libraries constructed using a Stranded mRNA Prep kit (Illumina), each from 500 ng RNA, as recommended. QC of libraries was performed using a Bioanalyzer 2100 and, following equimolar pooling of all 16 samples, RNA-seq was carried out on a NextSeq 2000 (Illumina) sequencer loaded at 750 pM. Fastq data was demultiplexed on completion, prior to downstream analysis.

The RNA-seq reads were pre-processed to remove adapters using Fastp (Chen *et al*., 2018). Low-quality reads, defined by a length < 30 or a quality score < 20, were eliminated. The processed reads were mapped to a high-quality potato reference transcriptome, which was assembled from a combination of RNA-seq short-reads and Iso-seq long-reads in a DM potato study (unpublished), using Salmon (Patro *et al*., 2017). The 3D RNA-seq App was applied to study the gene and transcript expression changes between wildtype (WT) and line 66 (L66), as well as samples exhibiting swelling and hooked phenotypes (Guo *et al*., 2022). Genes and transcripts with significant expression changes were determined with adjusted p-value < 0.05 and absolute log_2_-fold-change ≥ 1.

### Oxalate oxidase assay

Total oxalate oxidase activity was evaluated using an oxalate assay kit (Abcam ab241007) to quantify H_2_O_2_ production in the presence of oxalate following the manufacturer’s protocol. A total of 20 mg of fresh frozen leaf tissue was ground and resuspended in 200 µl ice-cold oxalate oxidase assay buffer then incubated for 10 minutes on ice. Following incubation, samples were centrifuged for 20 minutes at 10,000 g and 100 □l of the supernatant collected. Proteins were precipitated by addition of 200 µl 4.32 M ammonium sulphate followed by a 30 minute incubation on ice. Precipitated proteins were collected by centrifugation (10,000 g, 30 min), the supernatant discarded and protein resuspended in 100 □l fresh buffer to remove metabolites that may interfere with the assay. 40 µl of the reconstituted sample was then used in the final quantification assay.

### Construction of GERMIN-mRFP

The *GERMIN3* sequence was amplified from a template encoding GERMIN3 DMT46995-B3-B5 (Thermofisher Scientific) using attB adapter-flanked primers (GERMIN-FOR; 5’-AAAAAAGCAGGCTTCGAAGGAGATAGAACCATGGCTCTCAAGTACTTTGTATTAAC-3’ and GERMIN-REV; 5’-GGGGACCACTTTGTACAAGAAAGCTGGGTGGTTGTTATCCCACCAGAATTG-3’). The amplicon was recombined into pDONR207 using Gateway BP Clonase II (Thermofisher Scientific), then recombined into plasmid pK7RWG2 (Karimi *et al*., 2007) using Gateway LR Clonase II (Thermofisher Scientific).

### Live-Cell Imaging of Germin-mRFP

*Agrobacterium tumefaciens* cultures (strain AGL1) carrying the GERMIN3:RFP plasmid construct were prepared at OD600nm = 0.1 and infiltrated into the lower surface of *N. benthamiana* leaves expressing the ER marker GFP-ERD2 (Carette *et al*., 2000). For callose staining sample leaves were infiltrated with aniline blue stain 0.05 % in 0.067 M phosphate buffer, pH 8.5. Imaging of GERMIN3:RFP was performed on a Zeiss LSM 710 upright confocal laser scanning microscope (CLSM; Zeiss Jena, Germany) using a 40x objective, with an RFP excitation wavelength of 561 nm and emission collected at 590-630 nm. For aniline blue imaging an excitation wavelength of 405 nm was used and emission collected at 420 – 480 nm.

### Gating Experiment

*N. benthamiana* leaf epidermal cells were co-bombarded with mixtures of plasmid DNA by particle bombardment using a Handgun essentially as described by Gal-On *et al*. (1997). The plasmid mixtures comprised *35S::GFP:SPORAMIN* (Oparka *et al*., 1999) and *pK7RWG2::GERMIN3* (GERMIN3:RFP). Co-bombarded cells were examined by CLSM 2 days after bombardment. Cells were imaged using a 20× objective with both GFP and RFP imaged sequentially: GFP excitation at 488 nm, emission at 500 to 530 nm; RFP excitation at 561 nm, emission at 590 to 630 nm.

## Results

### Over-expression of GERMIN3 impacts meristem activation processes in potato

In order to probe the function of GERMIN3, transgenic lines were produced in *S. tuberosum* ssp. *andigena*, an obligate short-day potato (Jackson *et al*., 1996). The *GERMIN3* sequence used was from the Phureja double monoploid DM 1-3 515 R44 (based on transcript number PGSC0003DMT400046995 available at Spud DB (http://spuddb.uga.edu/)) and transgene expression was driven by a constitutive *35S* Cauliflower Mosaic Virus promoter. Twenty-six independent transgenic lines were screened for *GERMIN3* expression in leaves by qRT-PCR from tissue culture plantlets (Fig. **1a**). Sixteen lines exhibiting varying levels of *GERMIN3* expression were grown as described and tuber yield measured 47 days following sub-culturing. A strong and highly significant linear correlation between either leaf (Fig. **1b**) or stolon (Fig. **1c**) *GERMIN3* transcript abundance and tuber yield was observed. Five lines exhibiting high *GERMIN3* expression (lines 66, 69, 79, 89 and 91) were therefore selected for further analysis (Fig. **1a**). Although line 22 exhibited high abundance of *GERMIN3* transcripts, it grew poorly under tissue culture and glasshouse conditions and so was excluded from further analysis. In the selected lines, tuber yield was measured at two harvest points (47 and 93 days) with four out of the five overexpressing lines exhibiting higher yield than wild-type plants (Fig. **2a,c**; Fig. **S1**). Similarly, at the earlier time point the same four transgenic lines exhibited a higher tuber number (Fig. **2b**) although after 93 days only two lines maintained this phenotype (Fig. **2d**). These data suggest that over-expression of *GERMIN3* advances the date of tuber initiation and facilitates tuber bulking.

**Figure 1.**
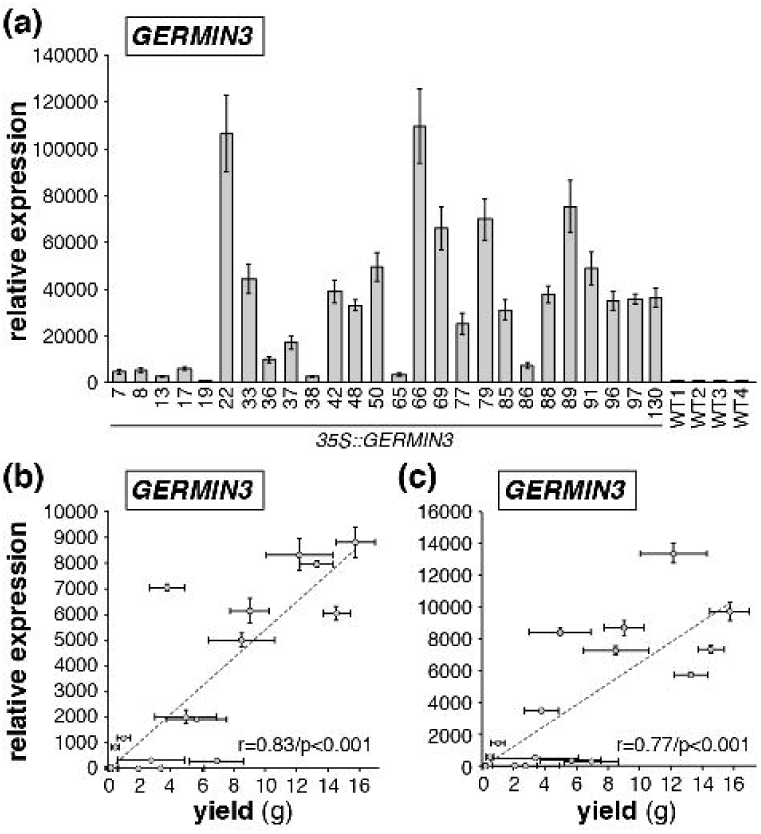
Relationship between GERMIN3 gene expression and tuber yield in GERMIN3 overexpressing S. tuberosum ssp. andigena transgenic lines. Relative abundance of the *GERMIN3* transcript in tissue culture leaflets in 26 overexpressing lines and Andigena controls (WT) are indicated (a) showing the variation in transcript abundance between lines. Sixteen lines were selected for further analysis and tuber yield was quantified 14 days after transfer to inductive short-day conditions and correlated against leaf (b) and stolon (c) *GERMIN3* transcript abundance. Expression levels were determined by RT-qPCR relative to the reference gene *StEF1□*. All data are represented as mean ± SE of 3 independent biological replicates. The correlation coefficient (r) and statistical significance (p) of the correlation are indicated in panels (b) and (c).

**Figure 2.**
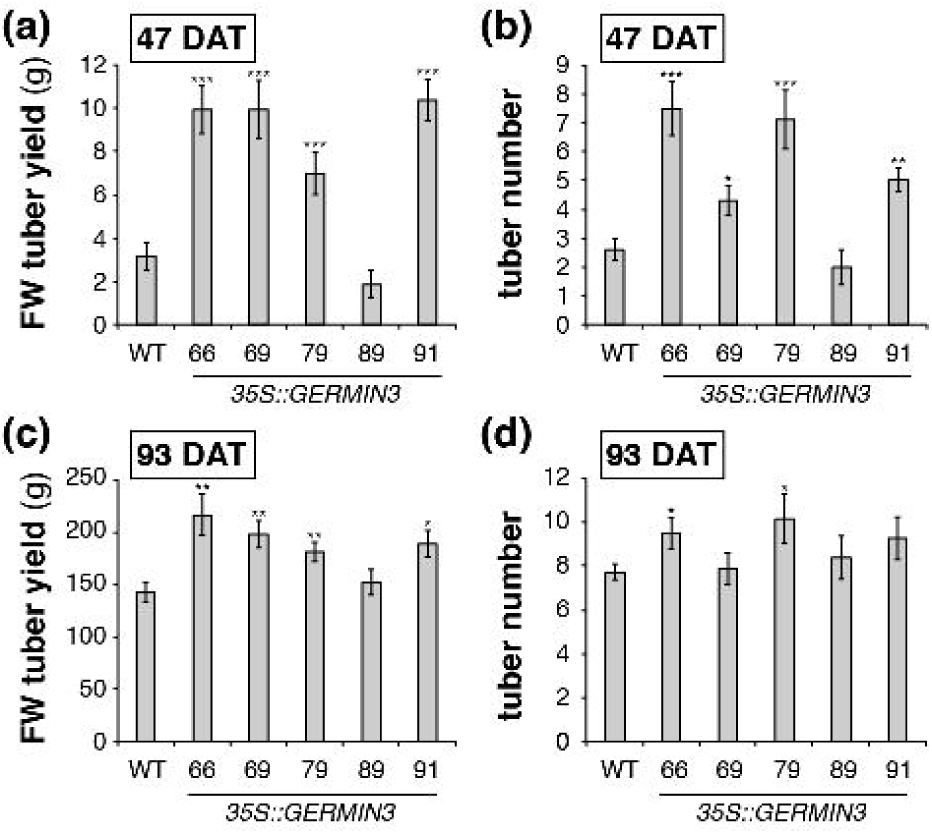
Tuber yield and number in wild-type and GERMIN3 over expressing S. tuberosum subsp. andigena transgenic lines. Tuber yield (FW) and number was measured in *35S::GERMIN3* lines and WT control. Plants were grown in pots under standard glasshouse conditions (18/15°C, 16 h daylength) for 35 days, then moved to a controlled environment (18/15°C) under short days (10 h daylength). Plants were harvested at 47 (a, b) and 93 (c, d) days after planting and tuber yield (a, c) and number (b, d) were recorded. All data are represented as means ± SE of 10 independent biological replicates at day 47 and 6 independent biological replicates at day 93. Asterisks denote values that were significantly different between transgenic lines and wild-type controls as determined by Student’s *t*-test (*p<0.05, **p<0.01, ***p<0.001).

A similar phenotype was observed in *S. tuberosum* cv. Desiree plants in which *GERMIN3* was expressed under 35S or the tuber-active *PATATIN* (*PAT*) promoter (Miroshnichenko *et al*., 2020) where two of three 35S and two of three *PAT* lines exhibited a significantly higher tuber yield 47 days after transfer to the glasshouse which was maintained in one 35S line and both *PAT* lines after 73 days (Fig. **S2**). On the contrary, knockdown of *GERMIN3* using RNAi technology had no impact on tuber yield despite a reduction in transcript abundance of up to 80% (Fig. **S2**).

The sprouting characteristics of tubers from the over-expressing lines were compared with those of WT plants. Sprouting was monitored in tubers stored at 10°C for up to 160 days. Whilst there was little difference in the time point when sprouting initiated between WT and transgenic lines (approximately 105 days), sprout number was significantly higher in four of the five *35S* lines relative to wild type indicating a loss of apical dominance (Fig. **3**).

**Figure 3.**
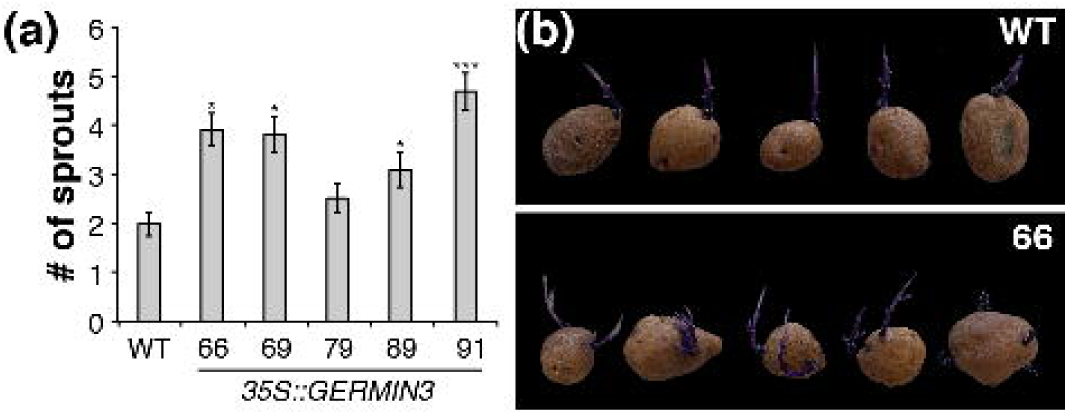
Number of sprouts recorded following tuber storage at 10°C in wild-type and GERMIN3 over-expressing S. tuberosum ssp. adigena lines. 20 tubers harvested from 10 replicate plants were stored at 10°C in the dark for 160 days and tuber dormancy was monitored. The number of individual tuber shoots with a length greater than 2 mm were counted and compared with ADG WT control (a). All data are represented as mean ± SE of 20 replicates (individual tubers). Asterisks denote values that were significantly different between transgenic lines and wild-type controls as determined by Student’s *t*-test (*p<0.05, ***p<0.001). The phenotypes of WT and over-expressing line 66 are indicated in panel (b).

The over-expressing lines also exhibited above-ground phenotypes, most notably a more highly branched growth pattern, in which the axillary buds, showing little or no outgrowth in wild-type plants, exhibited significant growth in the *35S* lines (Fig. **4**). The length of the stem derived from the axillary bud was measured in all nodes of the wild-type and overexpressing lines 66 and 79 (Fig. **4a**). Significant axillary bud outgrowth was observed particularly between node 6 and node 14. Internode length was also quantified indicating that all lines exhibited longer nodes in the middle region of the plant with overexpressing lines (Fig. **4b**), particularly line 66, having longer nodes than WT plants in many cases which led to an overall increase in plant height (Fig. **4c**). In addition to the axillary bud outgrowth in the OE lines that was not observed in WT (Fig. **4d-f**) we observed the growth of an organ resembling a stolon and an associated leaf in some nodes (Fig. **4g**). Taken together these data indicate that the reduced apical dominance observed in tubers which are modified subterranean stems is also apparent in aerial stems.

**Figure 4.**
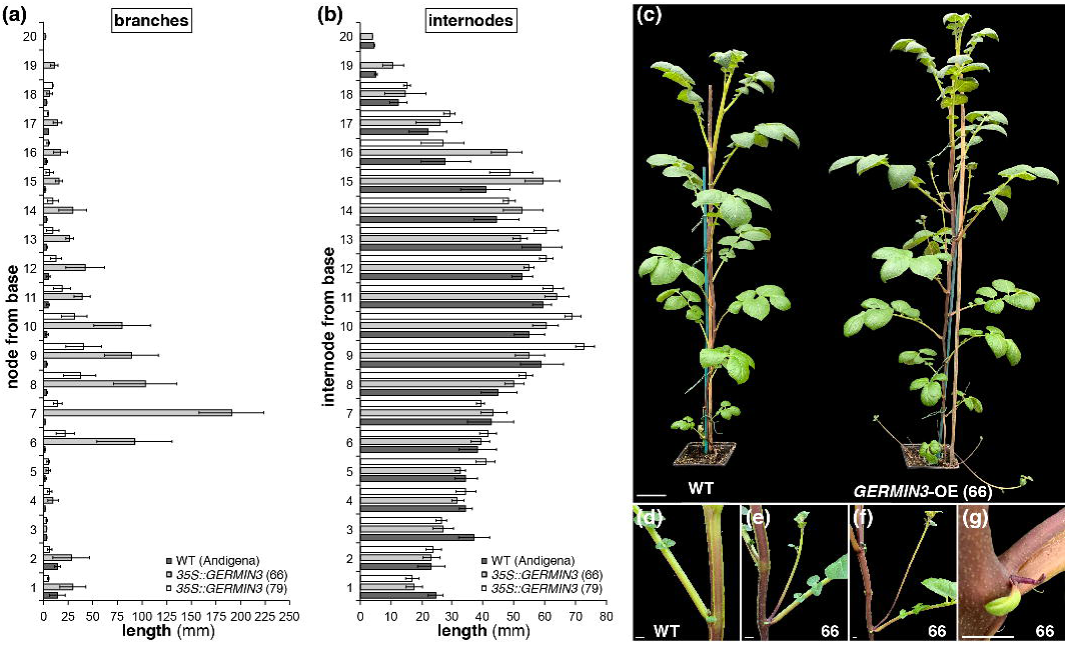
GERMIN3 over-expression results in increased stem axillary bud growth. GERMIN3 over-expressing lines exhibited stronger axillary bud growth than wild-type *S. tuberosum* ssp. *andigena* (a) and has altered stem length at some internodes (b). These phenotypes caused a more branched appearance and increased plant height (c). Axillary bud and internodal length was measured in 6 biological replicates 56 days after planting. All data are represented as mean ± SE.

Similar phenotypes were also observed in ‘Desiree’ plants expressing the *GERMIN3* construct under the *35S* promoter where axillary branches were typically more strongly developed (Fig. **S3a**), internode lengths were altered (Fig. **S3b**) and axillary bud outgrowth resulted in the production of stems, leaves and aerial stolons (Fig. **3c****-g**).

The flowering phenotype of overexpressing Andigena lines was also altered where andigena line 66 exhibited earlier flowering than wild-type lines after 56 days growth (Fig. **S4**).

### GERMIN3 is expressed in multiple organs of wild-type S. tuberosum ssp. andigena plants and is induced by tuberisation and starvation

The expression profile of *GERMIN3* in wild-type *S. tuberosum* ssp. *andigena* was determined by RT-qPCR analysis 14 days after transfer to inductive conditions when plants had stolons in multiple stages of development. A marked increase in transcript abundance was observed on transition from hooked to swelling stolons and expression remained elevated through late swelling and early tuber development (Fig. **5a**). On the contrary, where stolons were exposed to light following breaking through the surface of substrate, *GERMIN3* abundance was reduced.

**Figure 5.**
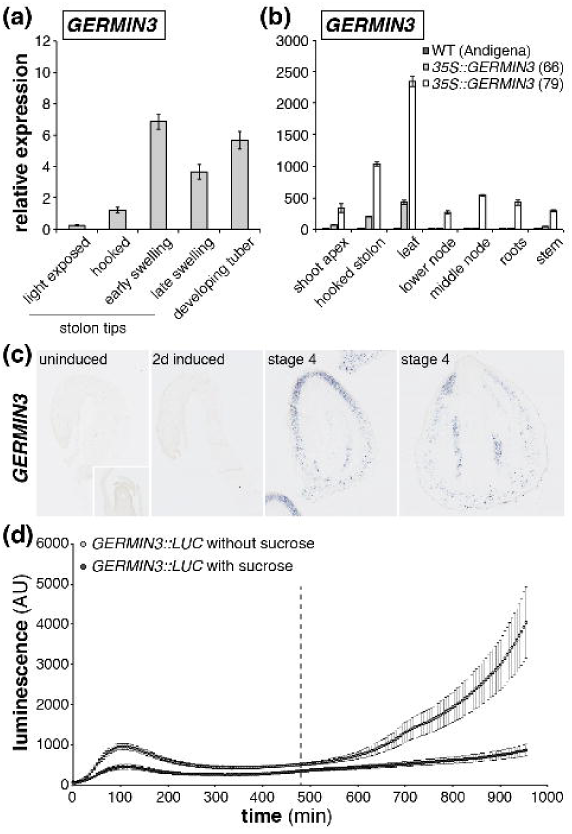
GERMIN3 transcripts are found in multiple organs of S. tuberosum ssp. andigena and are influenced by developmental and environmental cues. *GERMIN3* transcript abundance was quantified in stolon tips and developing tubers of wild-type *S. tuberosum* ssp. *andigena* (a) and in above and below ground organs of wild-type and two *GERMIN3* over-expressing lines (b). RNA *in situ* hybridisation was conducted in stolon tips and developing tubers of wild-type *S. tuberosum* ssp. *andigena GERMIN3*-specific antisense probes (c). The activity of the *GERMIN3* promoter was determined by transient expression of a *GERMIN::LUC* construct in *N. benthamiana* leaves (d). Leaf discs were prepared from the infiltrated leaves at the end of the day and promoter activity measured as luminescence following floating of discs on a buffer in the presence or absence of 2% sucrose. Dotted line indicates the end of the putative night.

Consistent with our observations regarding the aerial phenotype of *GERMIN3* overexpressing lines, the gene was also expressed in the shoot apex, the stem between nodes and nodal regions of plants as well as in the leaf and roots (Fig. **5b**). Expression of the gene under the *35S* promoter significantly enhanced transcript abundance in all tissues tested, particularly in line 79 (Fig. **5b**).

RNA *in situ* hybridisation experiments supported our observation that *GERMIN3* expression was limited in uninduced stolons where we were unable to detect any signal (Fig. **5c**). Similarly, no staining was observed with a full length *GERMIN3* antisense probe in 2 day induced stolons. On the contrary, stage 4 swellings exhibited significant staining associated with the cortex and vascular tissue supporting our RT-qPCR observations and indicating tissue-specific expression within the developing tuber.

To determine the upstream determinants of gene expression we conducted a series of promoter transactivation assays using group A bZIP transcription factors (StbZIP27, StbZIP36, StbZIP35, StbZIP66) to drive expression of a *GERMIN3::LUC* construct in transient assays. We failed to observe clear activation of expression by any of the bZIPs (data not shown). Unexpectedly, we observed an increase in luminescence in the absence of expression of any exogenous transcription factors after an extended period of measurement that coincided with the end of the putative night (Fig. **5d**). This increase in luminescence was abolished by inclusion of 2% sucrose in the buffer used to maintain leaf disks suggesting that *GERMIN3* transcription could be activated by endogenous transcriptional activators in response to starvation.

### Transcriptional profiles associated with altered GERMIN3 expression in stolons from transgenic lines

As tuberisation characteristics were altered in the *35S::GERMIN3* lines, we investigated the transcript level for the major tuberisation signal *SP6A* by RT-qPCR in leaves and stolons in wild-type plants and selected over-expressing lines. *SP6A* was detected in all leaf samples and no significant differences in abundance were observed between lines (Fig. **6a**). The *SP6A*-specific transcript was not detected in stolons prior to tuberisation except for line 66; however, transcripts were observed at similar abundances in swelling stolons of both WT and over-expressing lines (Fig. **6b**). Previously we have reported that TFL1B is an inhibitor of tuberisation that competes with SP6A for binding to FD and FDL in the tuberigen activation complex (Zhang *et al*., 2020). In leaves, *TFL1B* transcript abundances were similar in wild-type and over-expressing lines 66 and 79 although they were more abundant in leaves of *35S::GERMIN3* line 91 (Fig. **6c**). In hooked stolons significantly lower levels of *TFL1B* were detected in stolons prior to tuberisation in over-expressing lines than WT controls although abundance fell in all lines to a similar level following the initiation of stolon swelling (Fig. **6d**).

**Figure 6.**
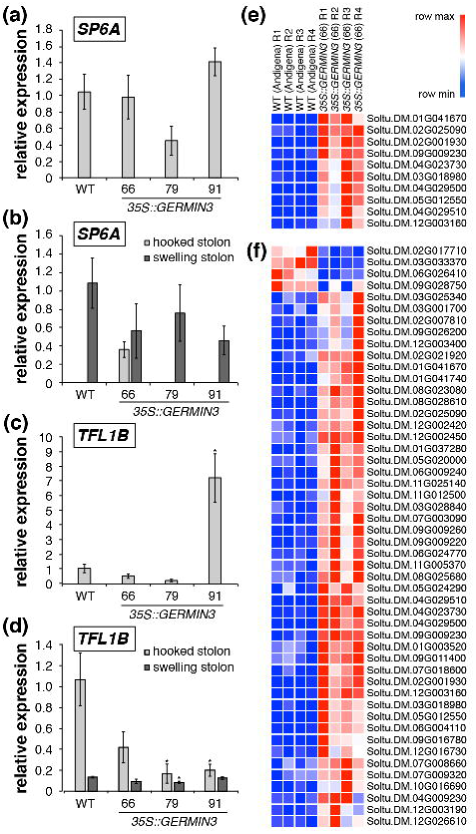
Abundance of key transcripts in stolons of wild type and GERMIN3 overexpressing lines. Abundance of *SP6A* (a, b) and *TFL1B* (c, d) transcripts were quantified by RT-qPCR in leaves (a, c) and stolons (b, d) in ADG *35S::GERMIN3* lines from plants cultivated for 47 days. Expression levels were determined relative to the reference gene *EF1*α. All data are represented as mean values ± SE of 3 independent biological replicates and asterisks denote values that were significantly different between transgenic lines and wild-type controls as determined by Student’s *t*-test (p<0.05). Transcripts that were significantly differentially abundant between wild type and overexpressing line 66 are also indicated for hooked (e) and swelling (f) stolons with each row showing relative abundance in individual replicates. Transcript identity is indicated to the right of each row and further details are provided in Table S3.

Transcriptomic analysis was performed in hooked and swelling stolons of wild-type and the transgenic *StGERMIN3* overexpressing line 66, selected for its high level of expression and consistent phenotype. Pairwise comparison of hooked and swelling stolons indicated that 5409 transcripts were significantly differentially abundant in wild-type plants with 2011 transcripts more abundant and 3398 transcripts less abundant in swelling than hooked stolons. In the GERMIN3 over expressing line 66, 3722 transcripts were differentially abundant with 1451 transcripts more abundant and 2271 less abundant in swelling stolons than in hooked stolons (Table **S2**). There was significant overlap in the identity of differentially abundant transcripts with a total of 3233 common transcripts in wild-type and over expressing line 66. Of the common transcripts, 1292 were more abundant in swelling stolons of both genotypes while 1941 were less abundant (Fig. **S5**).

This transcriptome analysis demonstrated an extensive level of changes upon tuber initiation in both the WT and *GERMIN3* over-expressing line 66. However, comparison of the two lines at identical stolon developmental stages revealed a relatively small number of differentially expressed genes (DEGs; Fig. **6e**). A total of 10 differentially abundant transcripts, all of which were more abundant in the *35S::GERMIN3* line relative to the WT line were present at the non-initiated hooked stolon stage. In swelling stolons, 50 transcripts exhibited differential abundance, 46 of which were more abundant and 4 less abundant in the *35S::GERMIN3* line relative to the WT line (Fig. **6f**). All DEG’s present in the hooked stage comparison were also present in the swelling stage gene list. A *GERMIN3* gene (Soltu.DM.01G041670) with a nucleotide similarity of 99.8% compared with the DM v4.03 transcript (PGSC0003DMT400046995) and used in the construction of the over-expressor construct presented in this manuscript was present in both gene lists. A second GLP gene (Soltu.DM.01G041740) was more highly expressed in the swelling stolon stage of the *35S::GERMIN3* line relative to the WT, the product of which may have a similar function to GERMIN3. Many of the remaining transcripts were poorly characterised although the list included cell-wall associated kinases (Soltu.DM.09G009260; Soltu.DM.09G009230 and Soltu.DM.09G009220) with a function in the regulation of cell expansion (Kohorn, 2016) as well as a transcript encoding an ADP-ribosylation factor GTPase (Soltu.DM.01G003520) that plays a role in cell division, expansion and cellulose production (Gebbie *et al*., 2005). Several transcripts encoding receptor-like kinases and a small number encoding transcription factors were also present within the DEG’s list as were a number of transcripts encoding cytochrome P450s and other oxidoreductases (Table **S3**).

Analysis of transcripts known to be associated with tuberisation such as *SP6A*, other PEPB’s and a range of transcripts associated with light signalling and circadian regulation indicated similar patterns of expression (Fig. **S6**). Taken together, these data indicate that molecular events associated with tuberisation were largely unaffected by overexpression of the *GERMIN3* gene.

### GERMIN3 over-expressing lines exhibit increased oxalate oxidase activity

Many GERMIN and GLP proteins exhibit oxalate oxidase activity (Lane *et al*., 1993; Sakamoto *et al*., 2015; Woo *et al*., 2000). To determine whether GERMIN3 has this activity, oxalate oxidase activity was quantified in extracts from leaves of WT and three highly expressing *35S::GERMIN3* lines. In extracts from WT leaves, 2.89 ± 0.32 □mol H_2_O_2_ min^-1^ mg FW^-1^ were generated in the presence of oxalate while the respective values for lines 66, 79 and 91 were 4.78 ± 0.56, 4.31 ± 0.52 and 5.49 ± 0.34 □mol H_2_O_2_ min^-1^ mg FW^-1^. Differences between *35S::GERMIN3* and WT lines were all highly significant (P<0.01, 0.05 and 0.001, respectively) according to the student’s *t*-Test with increases between 1.5- and 1.9-fold. These data suggest that GERMIN3 protein exhibiting oxalate oxidase activity accumulates in transgenic plants.

### GERMIN3 is localised in the Endoplasmic Reticulum and Impacts Plasmodesmatal Gating

The coding sequences of *GERMIN3* were expressed as C-terminal fusions to Red Fluorescent Protein (RFP). The constructs were introduced to source leaves of *N. benthamiana* expressing an ER-lumen localised GFP (Carette *et al*., 2000) by biolistic bombardment. Visualisation of the RFP-tagged GERMIN3 was used as a reporter for the presence of GERMIN3 which was localised by confocal microscopy. The protein clearly located to the endoplasmic reticulum where it colocalised with the GFP-tagged ER reporter (Fig. **7a-c**). Counterstaining with aniline blue highlighted the presence of callose associated with plasmodesmata at locations that were frequently coincident with the presence of ER-localised GERMIN3 (Fig. **7d-f**).

**Figure 7.**
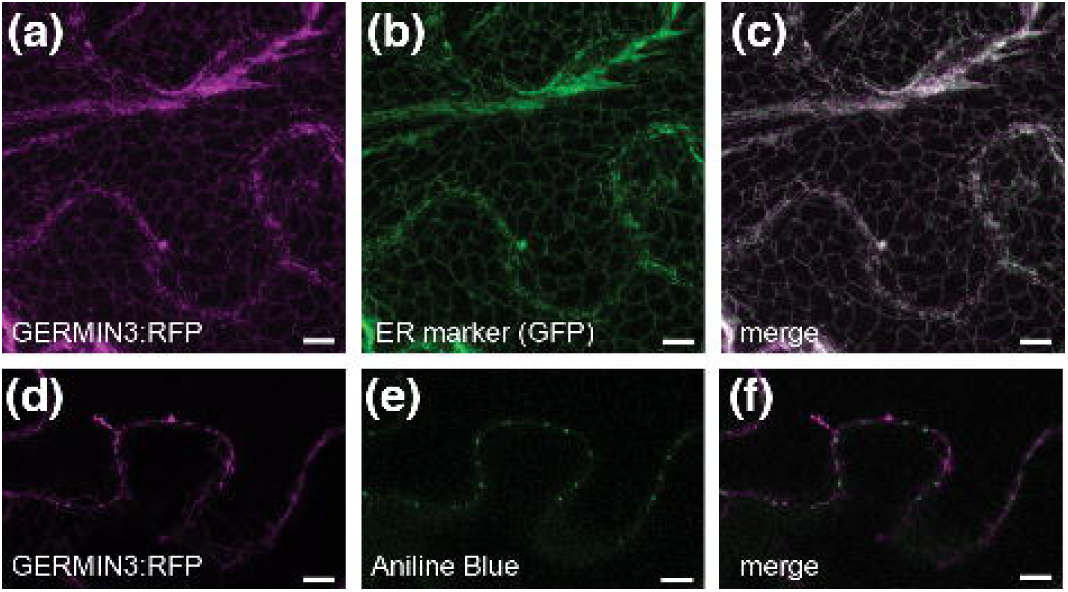
Intracellular localisation of GERMIN3:RFP. The GERMIN3:RFP fusion protein (a) localized to the endoplasmic reticulum as indicated by co-localisation with the GFP:ERD2 marker (b, c). A single stack of GERMIN3:RFP (d) indicates a close association with callose deposits at the neck of plasmodesmata stained with alanine blue (e, overlay in f). Scale bars represent 10 µm.

In order to determine whether GERMIN3 had any impact on plasmodesmal function, we used the plasmodesmatal gating assay described by Oparka *et al*. (1999). A plasmid expressing the sporamin-GFP fusion from a 35S promoter was bombarded into source leaves of *N. benthamiana* and the intercellular movement of the fusion protein was monitored by confocal microscopy. Other leaves were co-bombarded with the GFP:SPORAMIN construct and a construct expressing GERMIN3 tagged with RFP.

In control leaves that were bombarded only with the GFP:SPORAMIN construct, the GFP signal was primarily restricted to the bombarded cell with almost no trafficking to neighbouring cells (Table **S4**; Fig. **8b**). This result confirms previous work indicating that the low size exclusion limit of source leaf plasmodesmata restricts movement of the protein to adjacent cells (Oparka *et al*., 1999). In contrast, in cells that were co-bombarded and that expressed both the GERMIN3:RFP and GFP:SPORAMIN fusions, the frequency in which the SPORAMIN fusion was observed both in the bombarded cell and in neighbouring cells increased markedly (Table **S4**) while the GERMIN3 fusion was observed only in the bombarded cell (Fig. **8c,d**) suggesting that the presence of the GERMIN protein in the bombarded cell increased the size exclusion limit of plasmodesmata allowing passage of the GFP:SPORAMIN fusion between cells.

**Figure 8.**
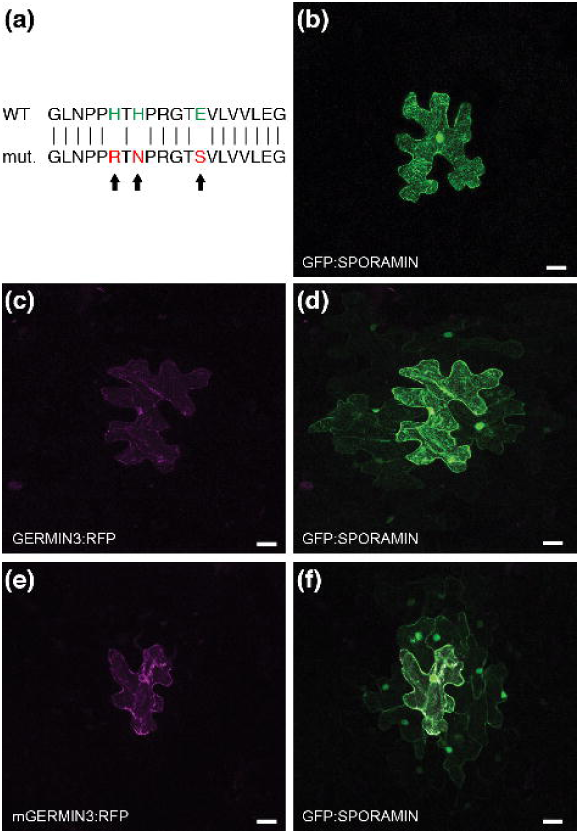
Plasmodesmal gating activity of wild-type and mutant Germin-RFP. *N. benthamiana* leaves were bombarded with plasmids encoding GFP:SPORAMIN alone (b) or co-bombarded with wild type GERMIN3:RFP and GFP:SPORAMIN (c, d). Alternatively, leaves were co-bombarded with a mutated GERMIN3:RFP (e) in which the Mn binding residues were changed to abolish binding (a) and the GFP:SPORAMIN construct (f). Scale bars represent 20 µm.

The results from the plasmodesmatal gating assays provide evidence for the function of GERMIN3. However, it is unknown whether the oxalate oxidase activity of the protein is necessary for its gating effect. The structure of many GERMIN and GLPs has been determined (eg da Cruz *et al*., 2019). The enzyme is a metalloprotein and binding of a Mn atom is required not only for oxalate oxidase but also for reported superoxide dismutase activity (Woo *et al*., 2000). The amino acid residues involved in Mn binding have been clearly identified (da Cruz *et al*., 2019; Sakamoto *et al*., 2015) and sequence alignment with potato GERMIN3 enable the corresponding residues (three histidine and a glutamic acid residue) to be identified in this protein. We synthesised a mutated GERMIN3 in which the Mn binding residues were changed (Fig. **8a**). We then used this mutated form in the gating assays where we found that the mutant protein exhibited a similar capacity to induce plasmodesmatal gating as the native protein (Table **S4**; Fig. **8e,f**). These data indicate that Mn-binding and oxalate oxidase/superoxidase dismutase activities are not required for plasmodesmatal gating.

## Discussion

### Regulation of the tuber life cycle

Tuber life cycle processes in potato are heavily regulated enabling the interface with environmental signalling to optimise and balance vegetative and reproductive growth and development. Significant progress has been made in recent years in our understanding of tuberisation signalling leading to a detailed model for tuberisation control (Kondhare *et al*., 2020; Zierer *et al*., 2021). As well as the FT orthologue, SP6A, other phloem mobile elements are important in tuberisation signalling (Hannapel *et al*., 2017). These include *BEL5* mRNA, the transcripts for other *BEL* family members (Ghate *et al*., 2017), and a number of small regulatory RNAs that impact on tuberisation signalling (Martin *et al*., 2009; Eviatar-Ribak *et al*., 2013; Lehretz *et al*., 2019).

These tuberisation signalling mechanisms converge on the tuberigen activation complex (TAC) that transcriptionally controls the cascade of gene expression that results in tuber formation. Our prior research identified *GERMIN3* as a target of TAC regulation and here we aimed to understand its detailed role in the tuber life-cycle and plant architecture.

### Over-expression of GERMIN3 accelerates meristem activation

The impact of constitutive *GERMIN3* expression on tuber development clearly demonstrated a positive effect on tuber yield, corresponding to the expression level in multiple independent transgenic lines. This was the case for batches of plants harvested 47 and 93 days after planting. These experiments were conducted in plants grown in pots and so may not reflect yields in the field, nevertheless there is clear potential for *GERMIN3* over-expression to have a positive impact under field conditions. In particular, the observation that *GERMIN3* acts downstream of the TAC provides an opportunity to maintain yields under conditions that impair TAC activation such as elevated temperature (Koch *et al*., 2024). Such an approach may be particularly beneficial if combined with transgenic manipulation of potato sink-source relationships using previously identified routes (Lehretz *et al*., 2021).

Over-expression of *GERMIN3* also resulted in other phenotypes associated with meristem activation processes. There was a significant effect on tuber sprouting with reduced apical dominance in the over-expressing lines resulting in the growth of more stems per tuber. Above-ground plant architecture was also impacted in *35S::GERMIN3* lines, with a significantly higher stem branched structure that indicated a loss of apical dominance.

### GERMIN3 controls PD gating supporting previous models of tuberisation/dormancy release

The study of Ham *et al*., 2012 provided evidence that several GLPs are located in the plasmodesmata where they have a role in the control of cell-to-cell trafficking (Ham *et al*., 2012). However, we are unaware of further work that extends these results. In this study, the GERMIN3:RFP fusion protein was clearly localised to the endoplasmic reticulum (ER). The ER is an essential component of plasmodesmata, the membrane-lined pores that interconnect plant cells. The desmotubule which traverses the centre of a plasmodesma is formed from, and continuous with, the cortical ER (Wright *et al*., 2006) and so our localisation studies are not inconsistent with a plasmodesmatal function.

Most GLP research implies the impact of these enzymes is due to their oxalate oxidase or superoxide dismutase activities. In the *35S::GERMIN3* lines generated in this study we measured increased oxalate oxidase activity. However, in view of the role of PD gating in meristem activation processes including tuber formation and sprouting (Viola *et al*., 2001; 2007), we wished to investigate whether GERMIN3 impacted on PD gating as observed in Arabidopsis. Using a well-established cell-to-cell trafficking assay we clearly demonstrated the impact of GERMIN3 expression, results that strongly suggest a role in the control of PD gating.

Previous studies have identified the germin amino acid ligands required for Mn binding and consequent oxalate oxidase and superoxide dismutase activities (Woo *et al*., 2000). We generated GERMIN3 mutated in these ligands and to abolish Mn binding and oxalate oxidase activity. The mutated form of the GERMIN3 retained activity in the cell-to-cell trafficking assay comparable to that of the wild-type GERMIN3. These data demonstrate that the oxalate oxidase activity of GERMIN3 is not required for the trafficking function. Further mutation studies will be required to identify the germin features that give rise to enhanced cell-to-cell trafficking. The experimental approach used in this study provides a tractable system for unravelling these effects. Detailed protein-protein interaction studies will shed further light on the precise mechanism of GERMIN3 activity.

### Concluding remarks

We propose that GERMIN3 and potentially other germin proteins may have a role in the control of plasmodesmatal gating, activating the switch from apoplastic to symplastic phloem unloading, marking the process of tuber-initiation and impacting on tuber sprouting in storage and above-ground stem branching. GERMIN3 acts downstream of the tuberigen activation complex and hence regulates tuberisation independently of environmental control. GERMIN3 is therefore a strong candidate for breeding and biotechnology for improved tuber yield in a wide variety of environments.

## Acknowledgements

This work was funded by the EU Horizon 2020 research and innovation programme ADAPT (Accelerated Development of Multiple Stress Tolerant Potato) grant agreement number GA 2020 862-858 and the Scottish Government Rural and Environmental Science and Analytical Services division as part of the strategic research programme 2022-2027.

## Author contributions

**Raymond Campbell**: Investigation, writing - original draft; **Graham Cowan**: Investigation; **Bernhard Wurzinger**: Investigation, writing - original draft; **Laurence Ducreaux**: Investigation, resources; **Jimmy Dessoly**: Investigation, resources; **Wenbin Guo**: Formal analysis; **Runxuan Zhang**: Formal analysis; **Jenny Morris**: Investigation; **Pete Hedley:** Data curation; **Vanessa Wahl**: Investigation, Resources, Supervision, Writing - original draft, Writing - Review & editing; **Mark Taylor**: Conceptualisation, Writing - original draft, Supervision, Project administration, Funding acquisition; **Robert Hancock**: Conceptualisation, Writing - original draft, Writing - Review & editing, Supervision, Project administration, Funding acquisition.

## Data availability

The data that supports the findings of this study are available in the supplementary material of this article.

