## Supplementary material for "GERMIN3 plays a role in plasmodesmatal gating to regulate meristem activation related to tuberisation, tuber dormancy release and stem branching in potato": Fig S1

Article acceptance date: Click here to enter a date.

The following Supporting Information is available for this article:

**Fig. S1** *Tuberisation phenotype of S. tuberosum ssp. andigena wild-type (a) and 35S::GERMIN3 line 66 (b).*

**Fig. S2** *Impact of altered expression of GERMIN3 on tuber yield in S. tuberosum cv. Desiree*

**Fig. S3** *Impact of altered expression of GERMIN3 on aerial phenotype of S. tuberosum cv. Desiree.*

**Fig. S4** *Flowering phenotype of S. tuberosum ssp. andigena wild-type and 35S::GERMIN3 line 66.*

**Fig. S5** *Common and unique differentially abundant transcripts in hooked and swelling stolons of wild-type and GERMIN3 over-expressing S. tuberosum ssp. andigena potato genotypes.*

***Fig. S6*** *Relative abundance of transcripts associated with tuberisation, light signalling and circadian regulation in stolons of wild-type and GERMIN3 over-expressing S. tuberosum ssp. andigena potato genotypes*.

**Table S1** *Primers and probes used for RT-qPCR*

**Table S2** *Relative abundance of transcripts in swelling compared with hooked stolons in S. tuberosum ssp. andigena (available as separate file)*

**Table S3** *Transcripts exhibiting significant differences in stolons between wild-type and GERMIN3 overexpressing line 66 genotypes (available as separate file)*

**Table S4** *GFP:SPORAMIN movement in N. benthamiana epidermal cells.*

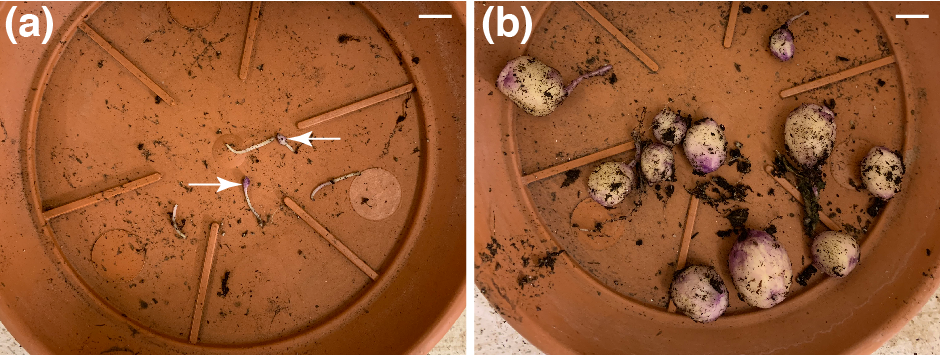

**Fig. S1** *Tuberisation phenotype of S. tuberosum ssp. andigena wild-type (a) and 35S::GERMIN3 line 66 (b).* Plants were grown as described and harvested after 47 days.  Stolons and tubers were carefully extracted from the substrate and photographed. Arrows in panel (a) indicate swelling stolons. Scale bars are 1 cm.

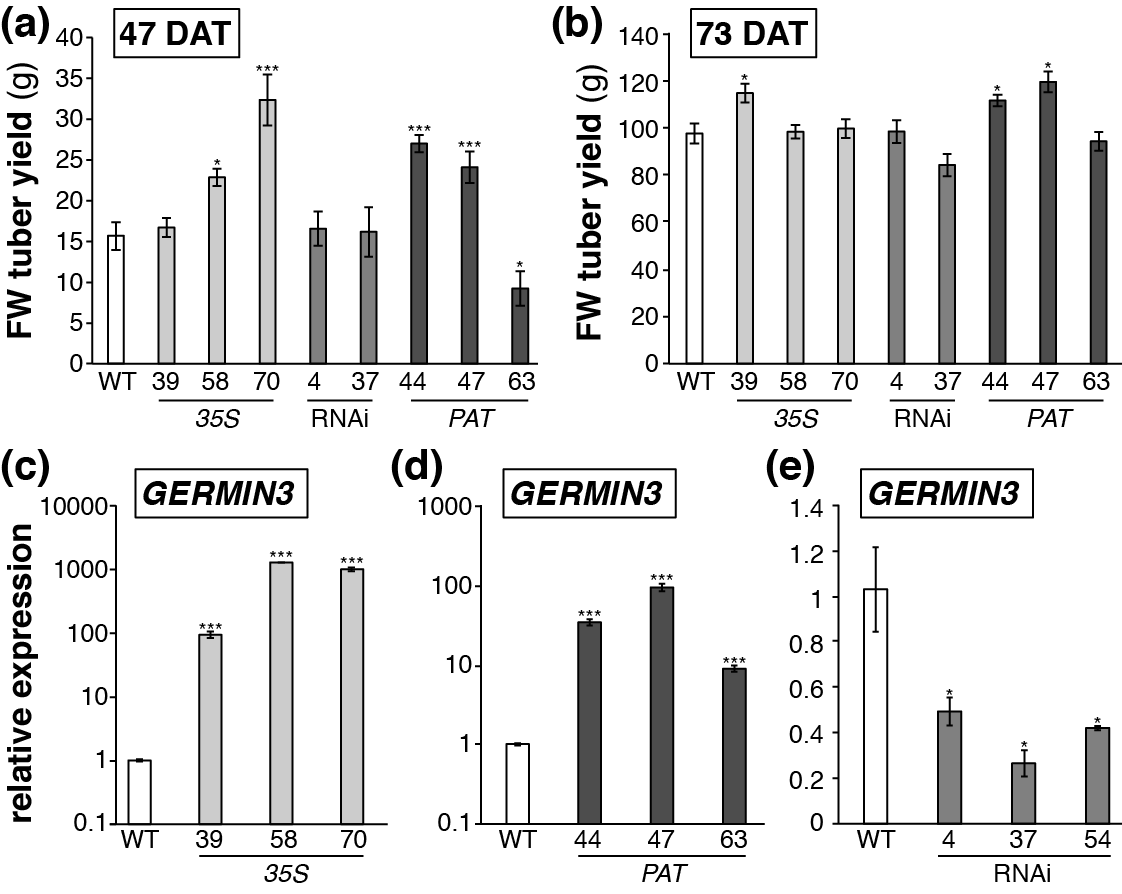

**Fig. S2** *Impact of altered expression of GERMIN3 on tuber yield in S. tuberosum cv. Desiree.* Wild type (WT), *35S::GERMIN3* (*35S*), *PATATIN::GERMIN3* (PAT) or *GERMIN3*-RNAi (RNAi) lines were transferred from tissue and grown for 47 (a) or 73 (b) days, respectively. Tubers were harvested and fresh weight recorded. *GERMIN3* transcript abundance was quantified by RT-qPCR in leaves of *35S* (c), developing tubers of *PAT* lines (d) and leaves of the RNAi lines (e). Expression levels were determined relative to the reference gene *EF1α*. All data are represented as mean values ± SE of 3 independent biological replicates and asterisks denote values that were significantly different between transgenic lines and wild-type controls as determined by Student’s *t*-test (p<0.05).

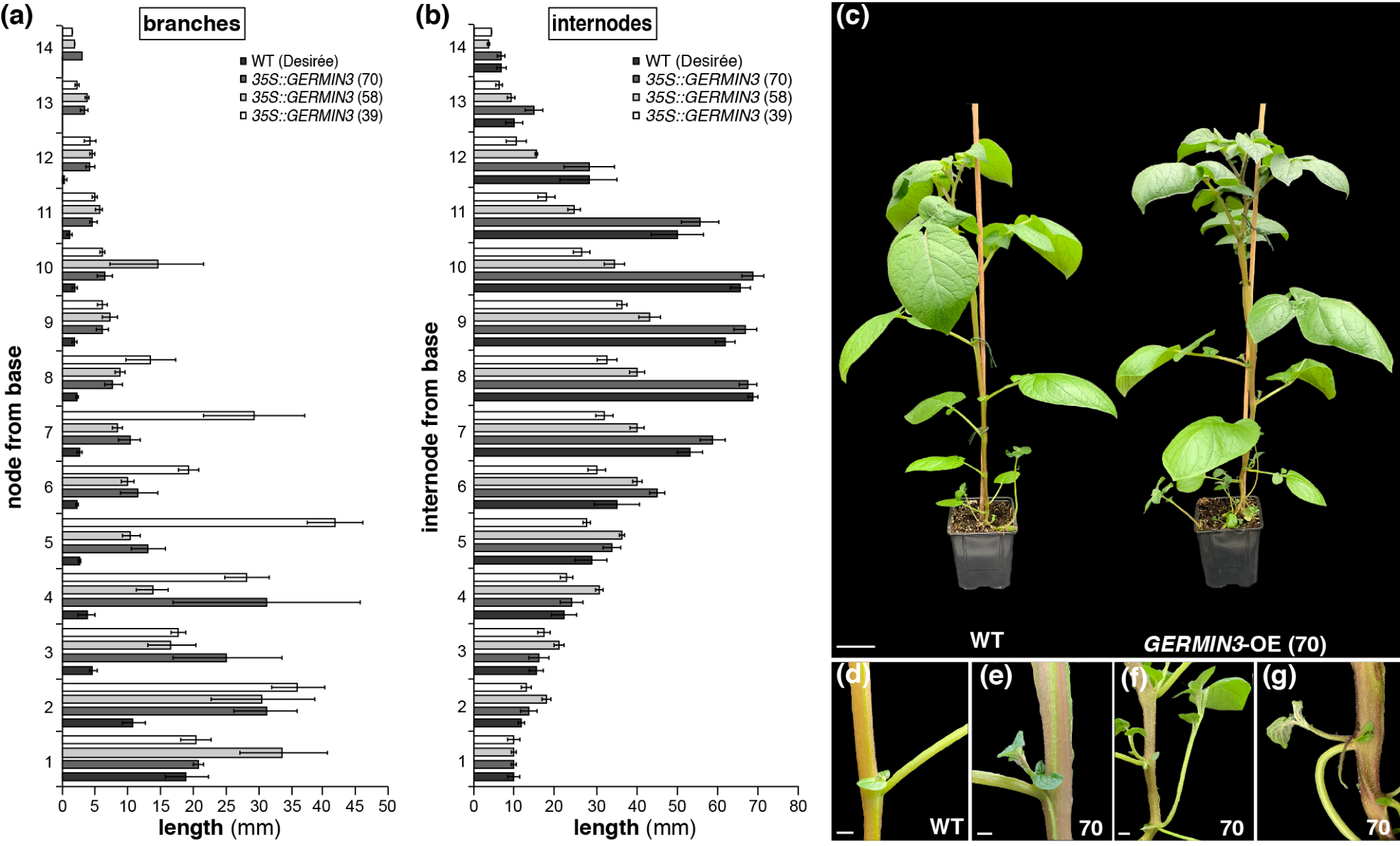

**Fig. S3** *Impact of altered expression of GERMIN3 on aerial phenotype of S. tuberosum cv. Desiree.* Over-expressing lines exhibited altered axillary bud growth (a) and internode length (b) relative to wild-type plants. This resulted in a more branched appearance (c) with the growth of stem-like and stolon-like structures that were reduced or absent in wild-type plants (d-g). Scale bars are 1cm.

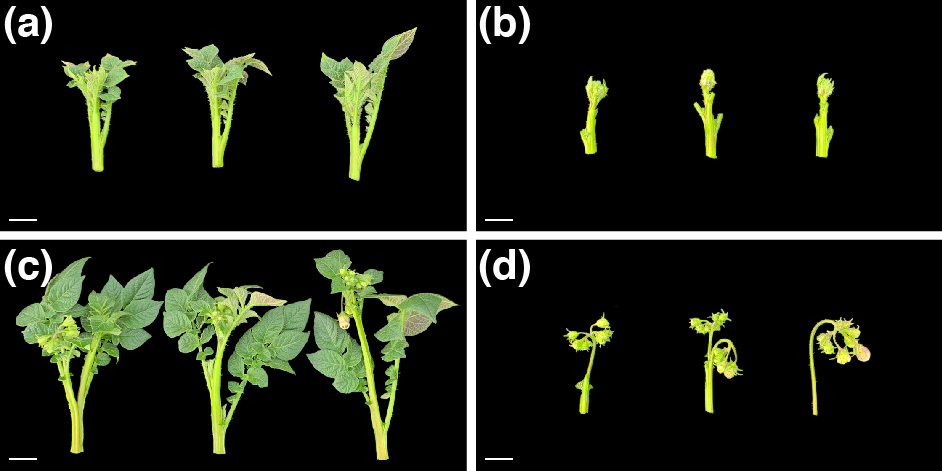

**Fig. S4** *Flowering phenotype of S. tuberosum ssp. andigena wild-type and 35S::GERMIN3 line 66.* Plants were grown under non-inductive conditions in the glasshouse under long days for 56 days. Apices from the wild type (a, b) and *35S::GERMIN3* line 66 (c, d) were removed and photographed whole (a, c) or following removal of leaves to reveal developing flowers (b, d). Scale bars are 1 cm.

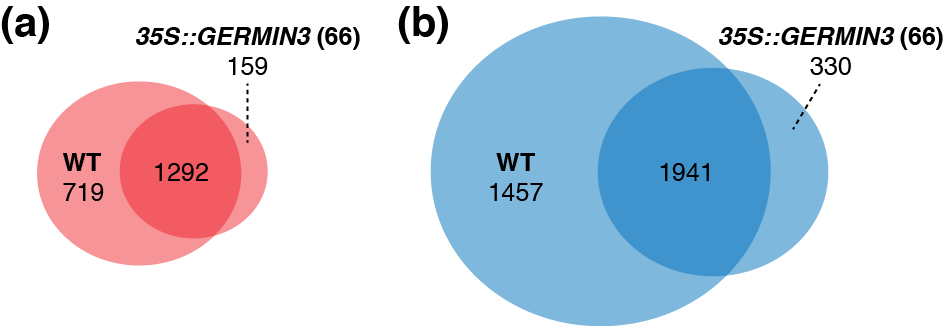

**Fig. S5** *Common and unique differentially abundant transcripts in hooked and swelling stolons of wild-type and GERMIN3 over expressing S. tuberosum ssp. andigena potato genotypes.* Venn diagrams indicate the number of unique and common transcripts that were significantly differentially abundant in swelling relative to hooked stolons in the wild type (WT) and *GERMIN3* overexpressing line 66. (a) Transcripts are more abundant in swelling relative to hooked stolons, (b) transcripts are less abundant in swelling relative to hooked stolons. Circles are drawn to scale relative to transcript number.

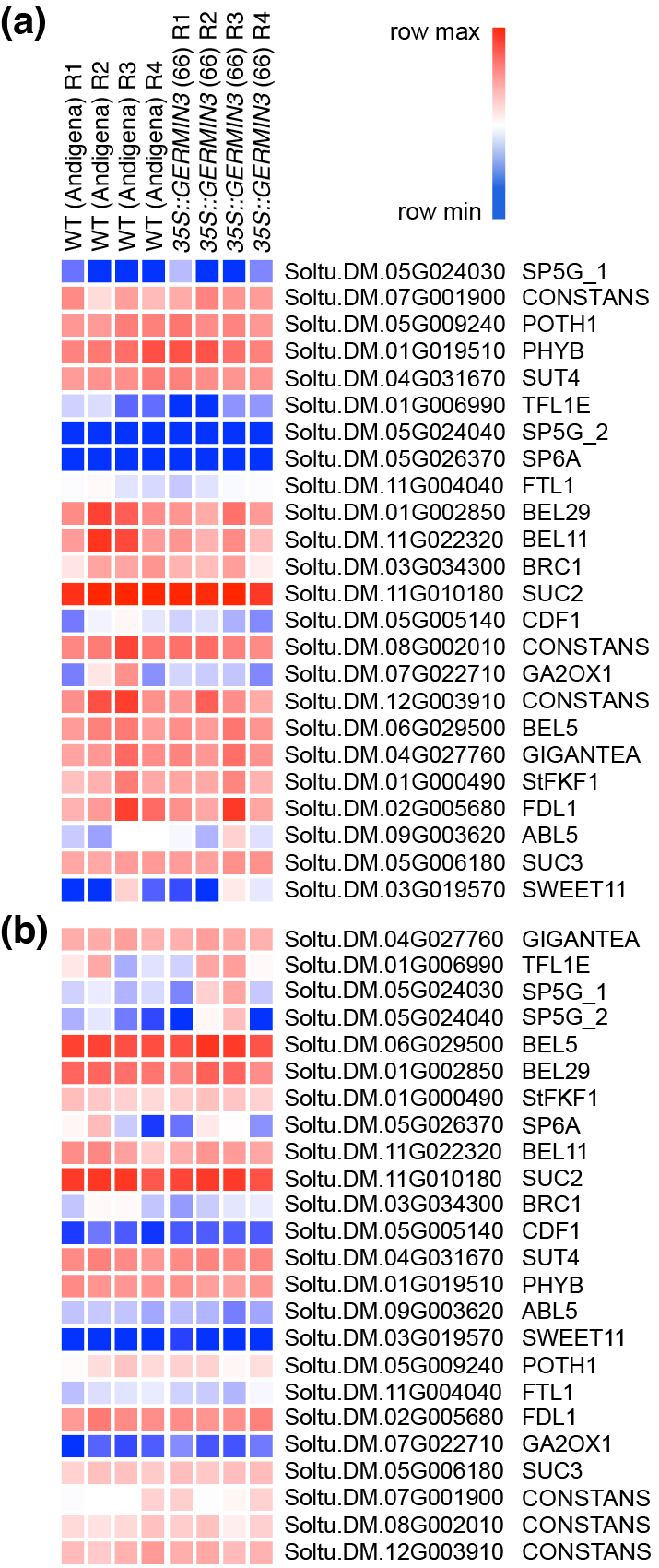

***Fig. S6*** *Relative abundance of transcripts associated with tuberisation, light signalling and circadian regulation in stolons of wild-type and GERMIN3 over-expressing S. tuberosum ssp. andigena potato genotypes.* Heat maps indicate the relative abundance of transcripts in hooked (a) or swelling (b) stolons of individual replicate plants. The accession and a brief transcript description is provided to the right of each row.

**Table S1.** *Primer probes for RT-qPCR*

| **Gene** | **PGSC ID** | **UPN** | **Forward** | **Reverse** |
| --- | --- | --- | --- | --- |
| **TFL1B** | Soltu.DM.03G017110 | 143 | AATGCCCAGAGAGAAACTGC | ATTTTGTGTGTGTGTGTGTCAAAT |
| **SP6A** | Soltu.DM.05G026370 | 138 | GGACGATCTTCGGAACTTTT | TCAAGTCAGGGTTGCTTG |
| **EF1**α | Soltu.DM.11G022490 | 117 | CTTGACGCTCTTGACCAGATT | GAAGACGGAGGGGTTTGTCT |

Gene name (Gene), potato genome sequencing consortium accession (PGSC ID) and sequences of primers used in PCR are indicated. UPN refers to the number of the Roche Universal Probe incorporate into the amplification reactions.

**Table S4** *SPORAMIN:GFP movement in N. benthamiana epidermal cells.*

| **Experiment** | **Construct** | | | | | |
| --- | --- | --- | --- | --- | --- | --- |
|  | GFP:SPORAMIN  - | | GFP:SPORAMIN  GERMIN3:RFP | | GFP:SPORAMIN  mGERMIN3:RFP | |
|  | Single cell | > 1 cell | Single cell | > 1 cell | Single cell | > 1 cell |
| **1** | 13 | 1 | 2 | 27 |  |  |
| **2** | 90 | 2 | 33 | 12 |  |  |
| **3** |  |  | 88 | 46 |  |  |
| **4** |  |  | 1 | 17 | 4 | 18 |

*N. benthamiana* leaves were bombarded with particles coated with constructs for the expression of GFP:SPORAMIN alone, GFP:SPORAMIN and GERMIN3:RFP or GFP:SPORAMIN and mGERMIN3:RFP carrying mutations in the Mn-binding site required for enzyme activity. Leaves were observed under a Zeiss LSM710 upright confocal laser scanning microscope (CLSM; Zeiss Jena, Germany) Using excitation wavelengths of 488 and 561 for GFP and RFP, respectively with corresponding emission collected at 500 – 530 for GFP and 590 – 630 nm for RFP. The table indicates the number of observation in which GFP fluorescence was restricted to a single cell or in which fluorescence was observed in more than one cell.
